# Semantic-Aware Energy-Efficient Operation in Smart Capsule Endoscopy

**DOI:** 10.64898/2026.03.17.712375

**Authors:** Mohammad Zoofaghari, Aliakbar Rahimifard, Saikat Chatterjee, Ilangko Balasingham

## Abstract

Goal-oriented semantic communication has recently emerged in wireless sensor-actuator networks, emphasizing the meaning and relevance of information over raw data delivery, thereby enabling resource-efficient telecommunication. This paradigm offers significant benefits for intra-body or implantable sensor-actuator networks, including dramatic reductions in bandwidth requirements, latency, and power consumption. In this paper, we address a patch-based energy-efficient anomaly detection method for smart capsule endoscopy. We propose a deep learningbased algorithm that employs the similarity between features extracted from measured images and a reference (normal) image as the detection metric. The algorithm is evaluated using a clinical dataset of capsule-captured images, combined with a simulated intra-body channel model. The results demonstrate that even with only 60% of the transmission power (relative to a standard link design for QPSK modulation) and 65% of the light intensity, the probability of anomaly detection remains above 85%, and it gradually improves as power and illumination levels increase. This improvement translates into a potential battery life extension of over 43%. The findings highlight the potential of semantic-aware, energy-efficient intra-body devices for more sustainable and effective medical interventions.

## I. Introduction

Resource-constrained Wireless Sensor-Actuator Networks (WSANs), are transforming domains such as digital healthcare, smart homes, industrial automation, environmental monitoring, and smart cities [1]. A typical WSAN consists of four key components: sensors, which detect and measure environmental parameters; actuators, which execute physical actions based on received commands; a control unit, which generates appropriate command signals; and a communication link, which enables information exchange between sensors, actuators, and the control unit, while also supporting remote monitoring and control.

A specialized subset of these systems, Wireless Body Sensor-Actuator Networks (WBSANs), integrates wearable sensors (e.g., activity trackers) or internal sensors (e.g., robotic wireless smart capsule endoscopy (SCE)) with therapeutic actuators to support real-time, adaptive interventions (Fig. 1) [2]. By tightly integrating sensing and actuation in closedloop systems, such as pancreatic control [3], pacemakers [4], brain–machine interfaces [5], and capsule endoscopy [6], WBSANs have significantly advanced the performance of intelligent healthcare systems. However, they require comprehensive control networks, especially when multiple sensing and actuation strategies must be coordinated to achieve a specific goal (e.g., adapting the system based on the detected activity class).

**Fig. 1.**
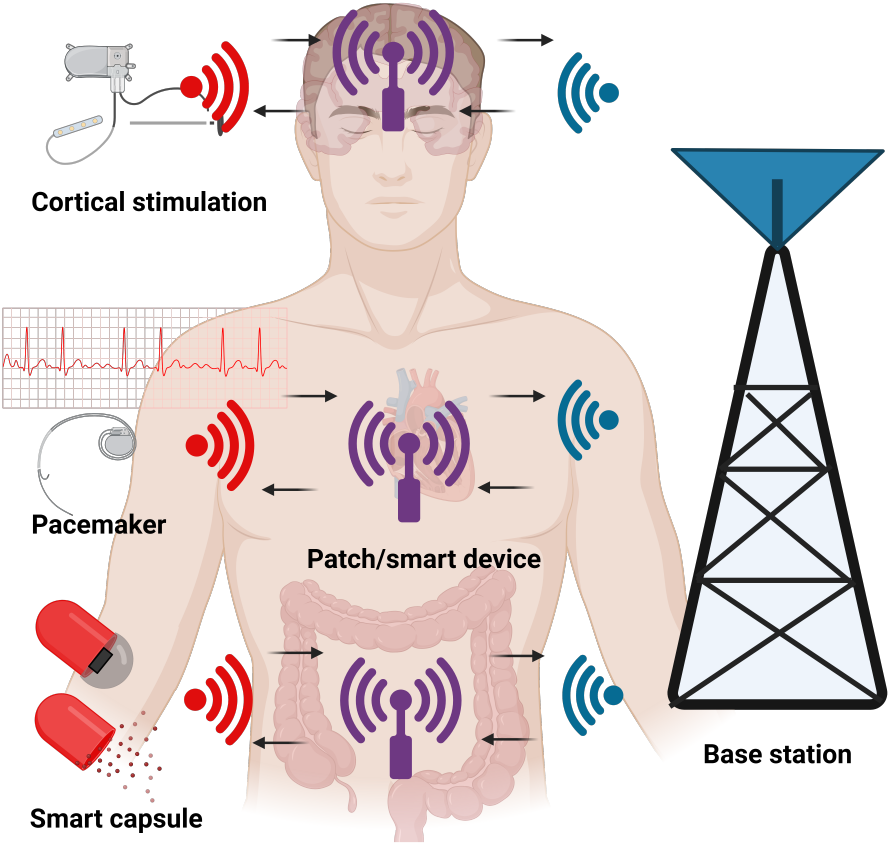
Closed-loop body area sensing actuation networks.

Challenges arise when communication delays between modules and the control unit lead to imprecise actuation, particularly in real-time medical applications (e.g., pacemakers, Fig. 1). Moreover, high-resolution sensing or frequent data transmission can exceed bandwidth limits in wireless systems. Intra-body communication is further constrained by lossy propagation environments, which degrade reliability and sensor performance. In addition, most remote sensors and actuators are battery-powered, and frequent communication intensifies power consumption. Uncontrolled actuation also risks excessive use of physicochemical resources.

One of the major shortcomings of current sensor-actuator systems lies in their neglect of the contextual factors that shape sensing and actuation. Data is often collected without consideration of the ultimate objective, while actuation is executed without accounting for its long-term effects [7]. This results in wasted resources and suboptimal system performance. For anomaly detection, frequent transmission of high-resolution images leads to wastage, as it leads to more information than necessary. Similarly, adjusting light intensity or triggering drug release without considering the actual disease state results in unnecessary energy consumption and wasted actuation resources. To address these limitations, semantic sensing, communication, and actuation have been proposed [8]. This paradigm ensures that only the most relevant data are sensed, transmitted, and acted upon, while actuation resources are applied only when they yield meaningful semantic effects.

In the gamut of WSAN, goal-oriented semantic communications integrate the semantic layer with the effectiveness layer, emphasizing the prioritization of transmitting semantic segments that are relevant to the system’s objectives [9]– [13]. This enables precise decision-making and successful achievement of the system’s ultimate goals. Within this framework, the notion of “meaning as importance” is introduced, whereby intelligent networked systems identify high-priority messages and dynamically allocate resources according to the importance attributed to the goal-driven value of each message. Goal-oriented sensing and actuation has been employed in many applications such as remote robot controllers and automated systems. For example, in [11], a prototype system was proposed in which an observer must transmit its sensory data to an executive agent responsible for carrying out a task (e.g., a mobile robot in a factory). The study then modeled the system as a partially observable Markov decision process (POMDP) and investigated the impact of incorporating semantic and effectiveness-oriented communication strategies on the overall system performance. Prior work has proposed transmitting semantic sensory data to control actuators, such as mobile robots in industrial settings [12], to minimize actuation costs and manage resource constraints by prioritizing actions [13].

Recently, semantic communication have been investigated in the context of WBASNs. For example, a context-aware semantic-based system for remote health monitoring has been developed and implemented, targeting groups such as elderly people with chronic diseases, pregnant women, and patients with physical disabilities [14]. In another example, [15] presents an innovative framework based on semantic communications for real-time electrocardiogram (ECG) signal monitoring. This framework operates on three levels: edge, relay, and cloud. At the edge level, a lightweight neural network identifies abnormal signals and forwards them to the relay where signals are semantically encoded and sent to the cloud for reconstruction and classification.

Most existing semantic-aware WSAN studies focus on communication resource allocation [16] rather than semantic-based energy adjustment of devices in physical systems. Similarly, semantic communication research in WBSAN focuses mainly on information transfer for tasks such as segmentation, classification, and diagnostic analysis, but does not provide a methodology or framework for energy-constrained resource allocation in intrabody devices.

In this work, for the first time, we present a goal-oriented approach to prolong the battery life of an intra-body device, demonstrated through a SCE dataset as a use case. We develop a deep learning-based method supporting end-to-end learning and feature representation [17] for anomaly detection. Our approach introduces a mechanism for event-aware adjustment of energy-intensive parameters in a closed-loop manner, guided by the semantic features. This goes beyond previous studies, which typically rely on reconstructed sensor data to guide actuation, by directly exploiting semantic features to optimize energy usage in real-time.

## II. Proposed Method

Intra-body communication faces strict constraints due to the highly attenuative and noisy channel. This emphasizes the need for goal-oriented power allocation to conserve battery life in the intra-body device while supporting reliable communication link design. Our approach addresses the anomaly detection objective in SCE, leveraging the semantic values of image light intensity and transmission power, enabling energy-efficient operation of the capsule. In SCE, the capsule is designed for non-invasive examination and treatment of the human digestive system by transmitting imaging data to an on-body smart device that uses expert knowledge to identify treatment-relevant tissue regions, polyps and abnormalities [18].

Fig.2 illustrates the proposed architecture for SCE anomaly detection, segmentation, and classification. The on-body device (patch) extracts key diagnostic images, such as those highlighting bleeding or polyps, and transmits them to an AI-edge system, enabling advanced tasks such as lesion segmentation, classification, and precise treatment recommendations (see Fig.2). Based on clinician-informed decisions, the edge device can direct the capsule to adjust its operation, such as modulating drug release, illumination, image resolution, frame rate, or localization strategy, to maximize diagnostic and therapeutic efficacy while minimizing off-target effects. Automatic patch-based adjustments could be applied to the capsule to enhance image quality in anomalous regions. This mechanism presents the remotely controlled smart capsule in a closed-loop manner as an emerging form of home-use medication.

**Fig. 2.**
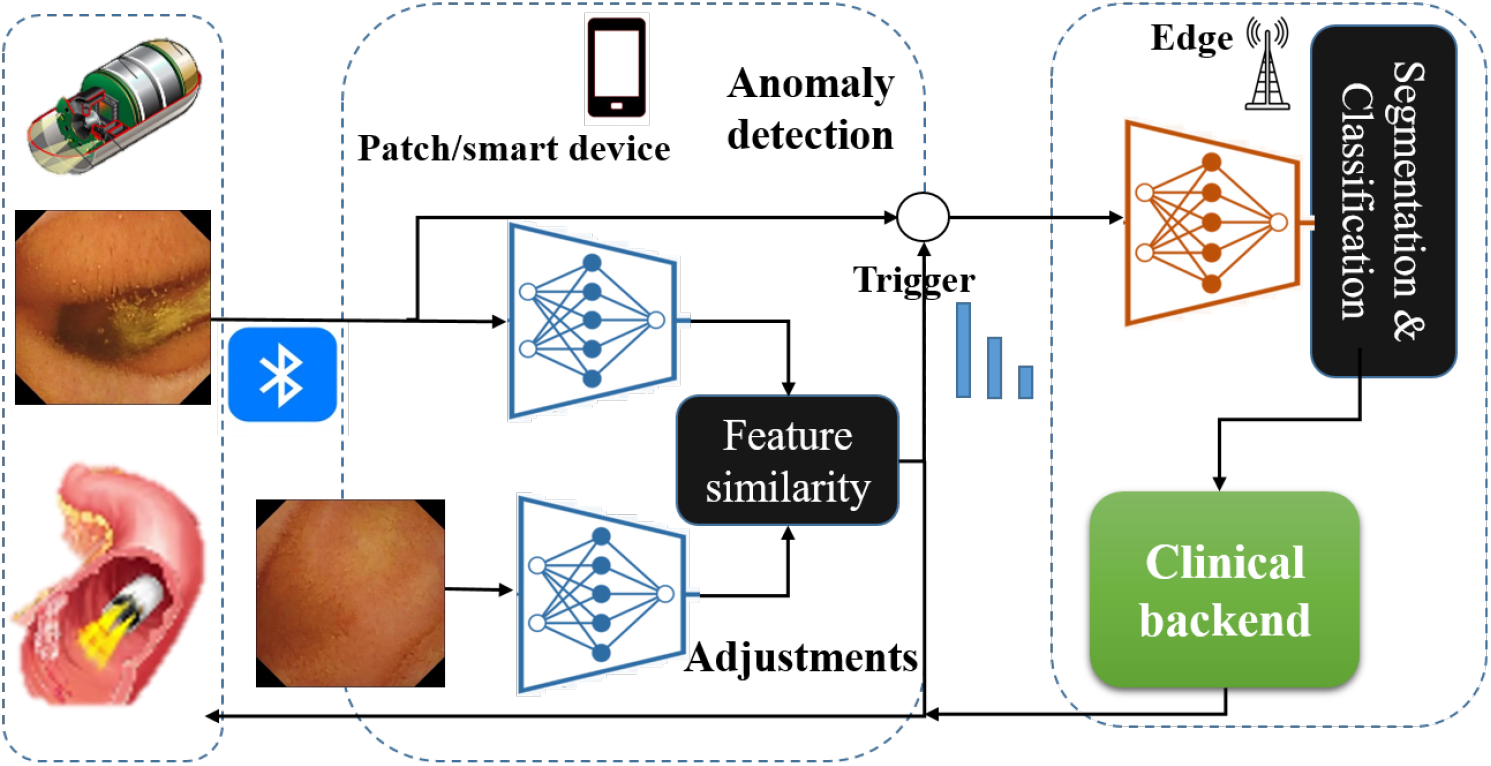
An architecture for the proposed deep learning-based anomaly detection and classification in smart capsule endoscopy (SCE).

A key factor in anomaly detection for SCE is the development and use of an appropriate metric. The metric should effectively distinguish between normal and anomalous tissue during scanning, while minimizing false alarms. We introduce the concept of *semantic similarity* for anomaly detection and SCE energy adjustment. Let *S*^*d*^ denote the desired signals representing target classes (Normal or Anomaly). Here, *S*^*d*^ may not directly correspond to labeled training samples (*S*^*l*^), but rather serve as ideal signal representations aligned with system objectives. In particular, *S*^*d*^ could be patient-specific and may represent the image of a lesion during the treatment process as the benchmark. We use a deep learing network as an encoder (ℰ ^(*L*)^(.)) to extract the features of the measured signals *S*^*m*^ and the desired signals *S*^*d*^. The corresponding features in the *L*^*th*^ layer are denoted as *F*^*m*^ and *F*^*d*^, and quantify semantic similarity of *S*^*m*^ and *S*^*d*^ as

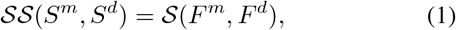

where *F*^*m*^ = ℰ^(*L*)^(*S*^*m*^) and *F*^*d*^ =ℰ ^(*L*)^(*S*^*d*^) and 𝒮 (·) denotes a similarity function selected from [19] (see Fig. 2). This formulation ensures that the similarity between the desired and measured images is captured semantically within the feature domain, serving as the proposed detection metric. Table I summarizes the steps of the algorithm for developing the semantic similarity function in terms of SCE energy-relevant parameters. This approach provides information beyond traditional similarity functions such as the Structural Similarity Index (SSIM), Learned Perceptual Image Patch Similarity (LPIPS), and Fréchet Inception Distance (FID), by extending the comparison from a syntactic level of images to a semantic level, thereby enabling more effective anomaly detection. Furthermore, it yields an analog measure of similarity between the measured image and the desired reference, moving beyond binary classification, and can be utilized for SCE parameter adjustment during diagnosis or treatment.

**TABLE 1.** Algorithmic representation of data flow and processing steps.

| Stage | Description / Operation |
| --- | --- |
| Input | $S^l = \{(S^l, y)\}$ : Labeled image data, $y \in \{\text{Normal, Anomaly}\}$<br>$S^m(I_L, \text{BER})$ : Measured data w.r.t. light intensity ($I_L$), and bit error rate (BER)<br>$S^d$ : Desired image<br>$L$ : Feature extraction layer, $S(\cdot)$ : Similarity function,<br>$\mathcal{E}$ : Encoder architecture with weights $\theta^e$ |
| Preprocessing | Apply noise reduction, filtering, and resampling. |
| Training Loop | 1. Encoder: $\mathcal{E}(S^l; \theta^e) \rightarrow z$<br>2. Predict: $\text{Class}_i(z) \rightarrow \hat{y}$<br>3. Optimize: $\arg \min_{\theta^e} \text{Loss}(y, \hat{y}) \rightarrow \theta_o^e$ |
| Feature Extraction | 4. Measured features: $\mathcal{E}^{(L)}(S^m; \theta_o^e) \rightarrow F^m$<br>5. Desired features: $\mathcal{E}^{(L)}(S^d; \theta_o^e) \rightarrow F^d$ |
| Output | 6. Compute performance: $S(F^m, F^d) \rightarrow SS(S^m, S^d)$ |

For anomaly detection, we set a threshold based on the Neyman–Pearson criterion and evaluate the performance of the process using the probability of detection and the probability of false alarm. This allows us to design the optimum threshold value for a given false alarm probability. We assume a Gaussian distribution of the semantic similarity 𝒮 𝒮 of the acquired images with the desired image belonging to the normal region. We apply maximum likelihood estimation to determine the standard deviations *σ*_*n*_, *σ*_*a*_ and the mean values *µ*_*n*_, *µ*_*a*_, corresponding to images from the normal and anomaly regions, respectively. Here, the probability of false alarm is given by

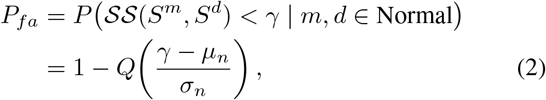

where *γ* is the decision threshold, *m* and *d* denote the measured and desired images, respectively, and *Q*(·) is the Q-function. Considering *P*_*fa*_ = 5%, the optimum threshold is

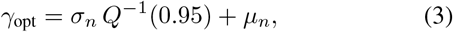

based on which the probability of anomaly detection is given by

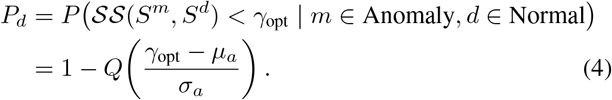

Please note that *σ*_*n*_, *σ*_*a*_, *µ*_*n*_, and *µ*_*a*_ are functions of the SCE parameters, such as light intensity *I*_*L*_ and bit error rate BER of transmitted images, as well as the deep encoder which provide semantic information to achieve a specific detection probability.

## III. Results and discussion

In this section, we present a numerical evaluation of patch-based anomaly detection using the proposed semantic similarity function, with respect to different capsule energy-related parameters. The challenges of edge-based segmentation and classification, combined with the EdgeNet communication will be addressed in future work.

We utilized the Kvasir-Capsule dataset, which contains more than 47, 000 clinical images across 14 anatomical and pathological classes [20], to train a convolutional neural network as encoder ℰ. To develop this network, we consider a binary classification setting in which all images of a specific lesion type are grouped into the anomaly class. We compare the performance of a pretrained lightweight network (SqueezeNet) and a pretrained moderate-weight network (GoogLeNet), both fine-tuned to our dataset through a transfer learning approach. SqueezeNet has very few parameters and low computational complexity, making it suitable for deployment on resource-constrained devices such as on-body patches. Its compactness, however, can limit representational capacity when dealing with complex patterns. In contrast, GoogLeNet is moderately heavier and computationally more demanding but offers richer feature extraction, which can enhance accuracy, particularly when processing low-quality images affected by energy and bandwidth constraints. Table II summarizes the performance of the models in the SCE dataset according to some typical metrics in deep learning [21]. As shown, although GoogLeNet has higher representational capacity, SqueezeNet achieves approximately 20% higher inference accuracy with more than 5M fewer parameters. This outcome is largely due to the relatively small dataset used in this study, where a network with fewer parameters fits the data more effectively.

**TABLE 2.** Comparative performance analysis of two CNN ARCHITECTURES ON A SMART CAPSULE ENDOSCOPY DATASET.

| Metric \ Model | SqueezeNet | GoogLeNet |
| --- | --- | --- |
| Inference accuracy (%) | 99.1 | 80.0 |
| F1-score (%) | 95.0 | 78.7 |
| Recall (%) | 93.1 | 76.5 |
| Precision (%) | 98.2 | 79.2 |
| No. of parameters (millions) | 1.24 | 7 |

To evaluate the performance of two models on video frames of a SCE clip streamed to the patch, Fig. 3 illustrates the semantic similarity between patch-received frames and a reference image from the normal class. We employ the cosine similarity function 𝒮, applied to both the pre-softmax features and the softmax output, denoted as *CSS*. As illustrated, *CCS* based on the softmax output clearly highlights the region annotated as erosion by the clinician for both models. However, pre-softmax-based *CSS* exhibits fewer false alarms, albeit with lower *CSS* drop within the anomaly region and higher fluctuation in normal region, suggesting use of an optimal threshold for a specific false alarm probability. To assess model performance under light-adjusted SCE conditions, Fig. 4 presents the *CSS* when the light intensity of the measured frames is reduced to 80%. As shown, SqueezeNet fails to differentiate the anomaly, whereas GoogLeNet exhibits a clear *CSS* drop in both the pre-softmax features and the softmax output, underscoring its robustness under light-limited conditions. A similar effect can also be observed for images constrained by the transmission power limitations.

**Fig. 3.**
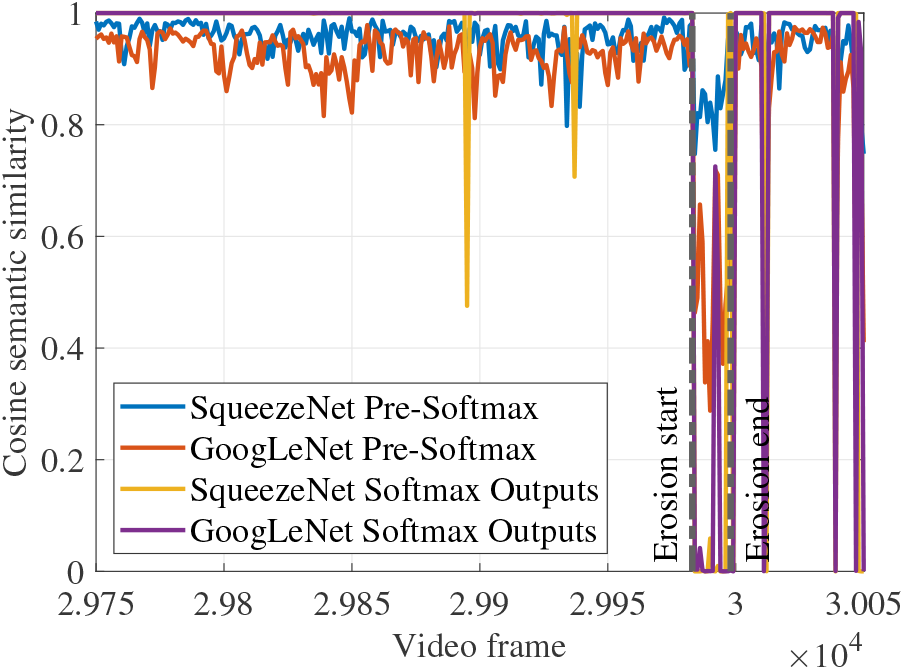
Semantic similarity of video frames captured by smart capsule endoscopy with a desired image in the normal class for two feature sets and models.

**Fig. 4.**
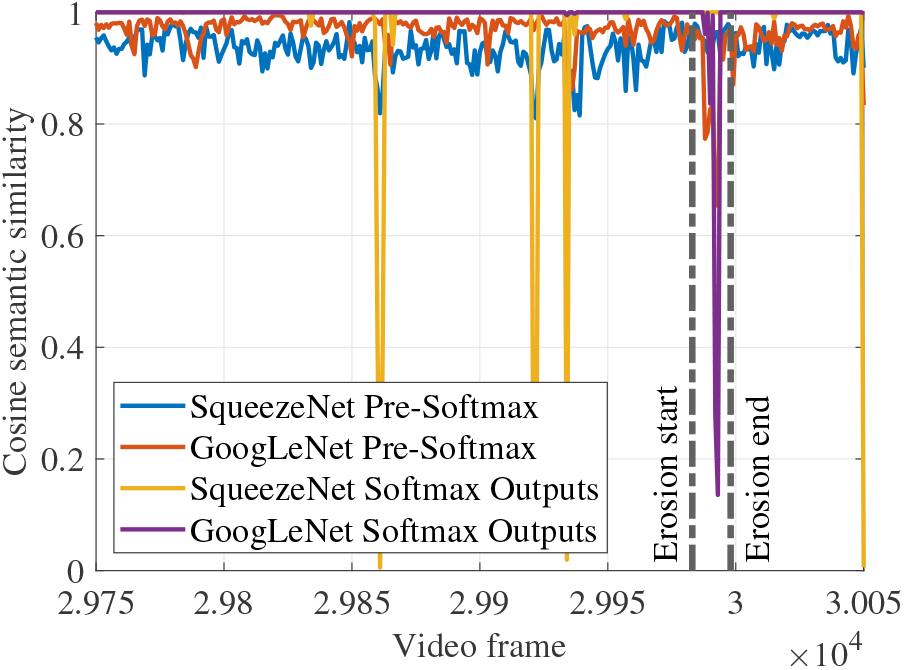
Semantic similarity of video frames captured by smart capsule endoscopy with a desired image in the normal class for two feature sets and models at light = 80%.

To evaluate the performance of *CSS* given by GoogLeNet under different lighting conditions, Fig 5 shows the *CSS* values of pre-softmax features and softmax output for 100 light-adjusted measured images considering a desired full-light image from the normal class, plotted as a function of normalized light intensity *I*_*L*_. As observed, *CSS* reaches saturation for light intensities above 70%, with only a small variation range. In contrast, at light intensities below 70%, *CSS* derived from softmax output exhibits a very large variation range, reducing the reliability of these features for anomaly detection. However, the variance of *CSS* remains low when based on pre-softmax features, clearly indicating that these features are more robust to illumination changes and therefore more suitable for real-time, low-light anomaly detection.

**Fig. 5.**
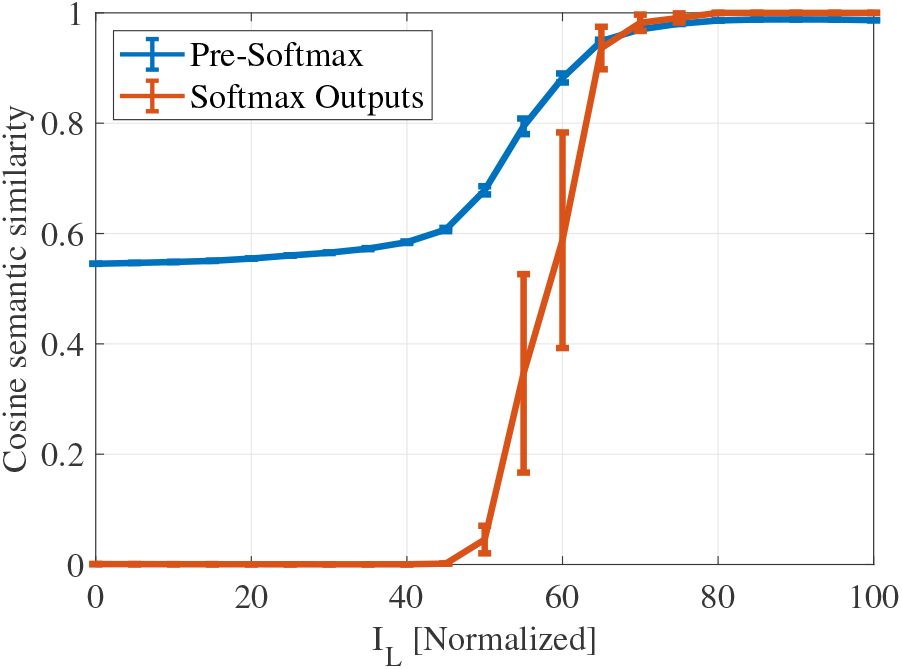
Average values and variation ranges of CSS as a function of light intensity, computed over 100 normal-class images using the GoogLeNet model.

In order to investigate the joint effect of BER and *I*_*L*_ on the optimum threshold value and the probability of anomaly detection, we generated Fig. 6 and Fig. 7 for pre-softmax features and softmax output, respectively.

**Fig. 6.**
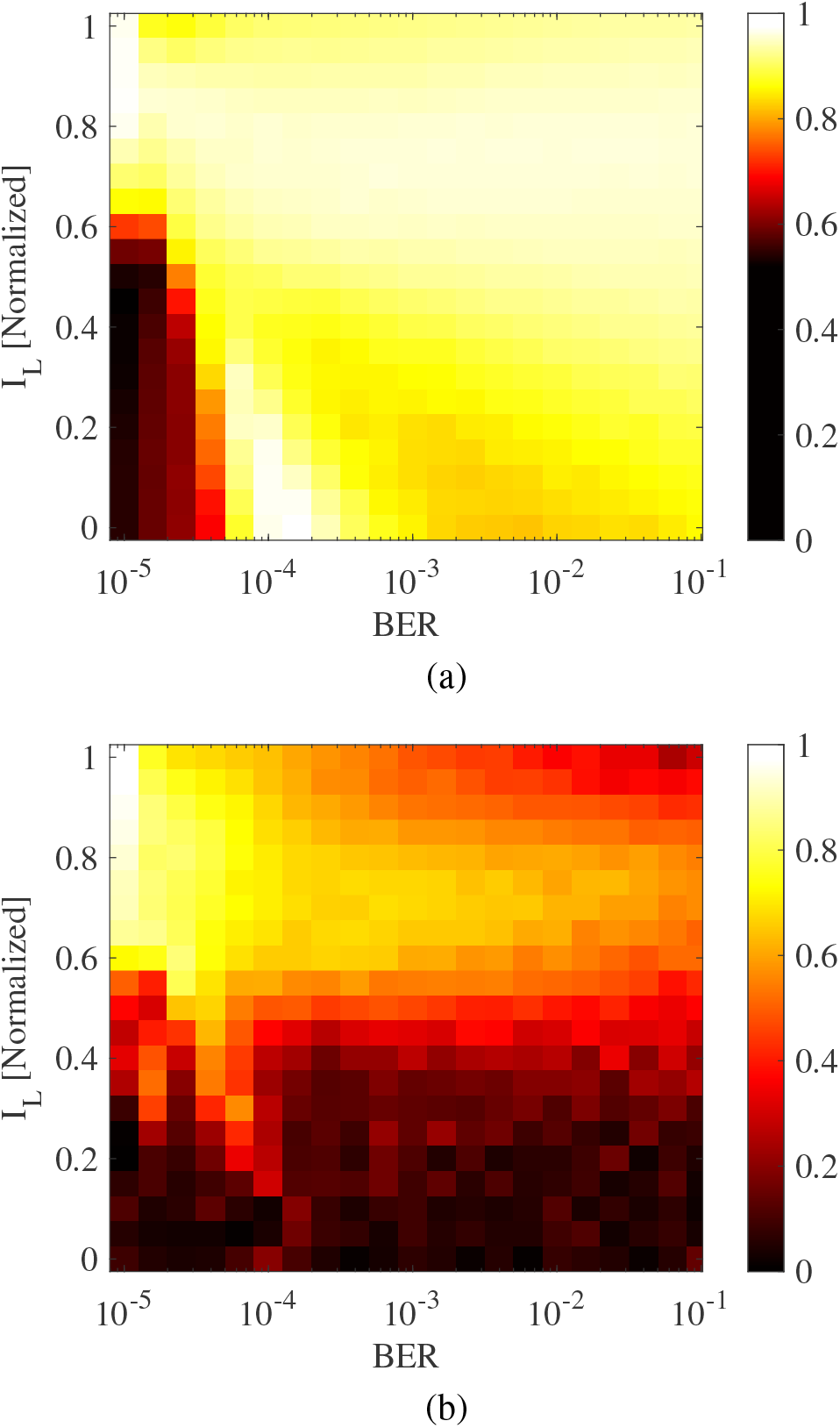
(a) Optimum threshold value *γ*_opt_ and (b) detection probability *P*_*d*_ in terms of normalized light intensity and BER for the GoogLeNet encoder and pre-softmax features.

**Fig. 7.**
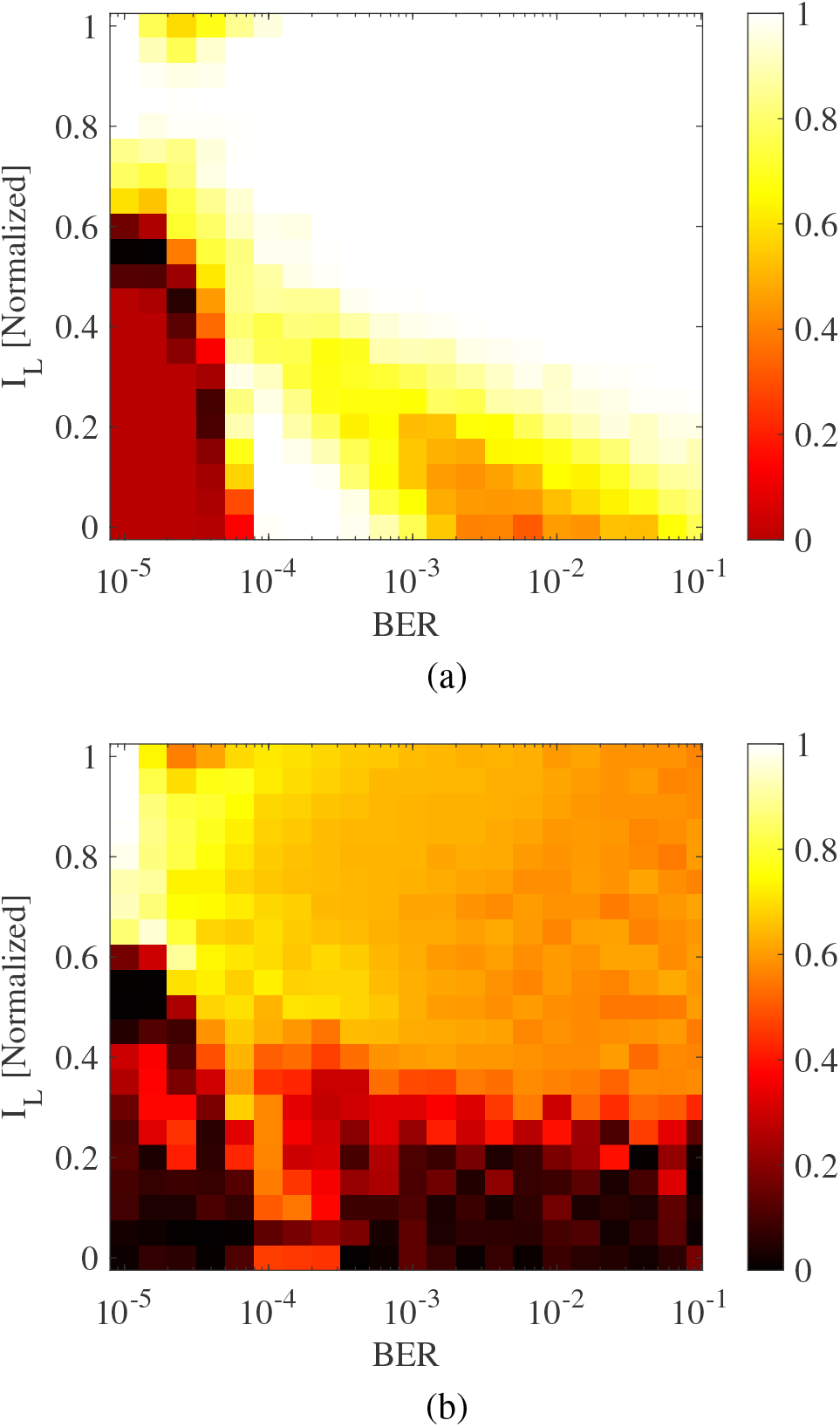
(a) Optimum threshold value *γ*_opt_ and (b) detection probability *P*_*d*_ in terms of normalized light intensity and BER for the GoogLeNet encoder and softmax output.

We jointly adjusted BER and *I*_*L*_ for 100 samples measured from normal and anomaly class images, and calculated *σ*_*n*_, *σ*_*a*_, *µ*_*n*_, and *µ*_*a*_ for the obtained *CSS* values based on the proposed method. These values were then used in (3) and (4) to yield *γ*_opt_ and *P*_*d*_, respectively. As shown in Fig. 6a and Fig. 7a, the optimal threshold *γ*_opt_ suggests a lower threshold for low-energy SCE to avoid a high false alarm rate, as there is higher fluctuation of *CSS* under low power conditions. The detection probability results indicate that, for BER ≤2.5 × 10^−5^ and 0.65 ≤*I*_*L*_ in the case of pre-softmax features, and for BER ≤ 2.5 × 10^−5^ and 0.8 ≤ *I*_*L*_ in the case of softmax output, we obtain *P*_*d*_ ≈ 0.85. Given a SCE communication link with standard AWGN QPSK modulation, the normalized transmit power ratio is expressed as

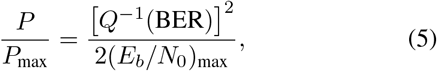

where (*E*_*b*_*/N*_0_)_max_ denotes the reliable bit energy-to-noise ratio, assumed here to be 11.3 dB for BER = 10^−7^. Taking into account the threshold values for BER, this would result in a reduction in the transmission power of around 40%. Given the assumed percentages of light and power reduction, and assuming that these parameters account for 40% of the SCE battery power, the battery life can be improved by approximately 43% for pre-softmax features and around 32% for softmax output. This demonstrates a substantial extension of battery life, particularly when exploiting the pre-softmax features that incorporate semantic information about light and transmission power.

## IV. Conclusion

This paper investigates a semantic approach to anomaly detection in wireless body area networks, demonstrated as a proof of concept for smart capsule endoscopy. The proposed technique facilitates event-aware data transmission from an on-body device to the edge, enabling computationally intensive segmentation and classification of SCE images and save the edge communication resources. The focus of this work is on prolonging capsule battery life by introducing semantic values for LED light intensity and wireless transmission power to achieve specific anomaly detection and false alarm probabilities. This approach benefits the intra-body communication link design, which faces strict constraints on the SNR due to the highly attenuative and noisy environment. Furthermore, our approach paves the way for a novel strategy of event-aware SCE parameter adjustment, such as increasing light intensity in regions of anomalies to improve segmentation and classification performance. Future studies are needed to identify the optimal network architecture, taking into account capsule resource constraints and the feasibility of deploying the model on an on-body device. Additionally, the concept of predictive control could be applied to enable automatic closedloop operation using the proposed semantic similarity function.

